# Curing of a field strain of *Salmonella enterica* serovar Infantis isolated from poultry from its highly stable pESI like plasmid

**DOI:** 10.1101/2024.02.01.578384

**Authors:** Nadia Gruzdev, Chen Katz, Itamar Yadid

**Affiliations:** Migal-Galilee Research Institute, Kiryat-Shmona 1101602, Israel; Tel-Hai College, Upper Galilee, 1220800, Israel

**Keywords:** *Salmonella enterica* serovar Infantis, Megaplasmid-pESI, plasmid curing, toxinantitoxin addiction systems

## Abstract

*Salmonella enterica* serovar Infantis (*S*. Infantis) is an important emerging pathogen, associated with poultry and poultry products and related to an increasing number of human infections in many countries. A concerning trend among *S*. Infantis isolates is the presence of plasmid-mediated multi-drug resistance. In many instances, the genes responsible for this resistance are carried on a megaplasmid known as the plasmid of emerging *S*. Infantis (pESI) or pESI like plasmids. Plasmids can be remarkably stable due to the presence of multiple replicons and post-segregational killing systems (PSKs), which contribute to their maintenance within bacterial populations. To enhance our understanding of *S*. Infantis and its multidrug resistance determinants toward the development of new vaccination strategies, we have devised a method for targeted plasmid curing. This approach effectively overcomes plasmid addiction by leveraging the temporal overproduction of specific antitoxins coupled with the deletion of the partition region. By employing this strategy, we successfully generated a plasmid-free strain from a field isolate derived from *S*. Infantis 119944.

This method provides valuable tools for studying *S*. Infantis and its plasmid-borne multidrug resistance mechanisms.

## 1. Introduction

*Salmonella enterica* is a human and animal bacterial pathogen responsible for global pandemics of food-borne infections. The species *S. enterica* is classified into six subspecies and includes over 2600 serovars [1]. Non-typhoid *Salmonella* (NTS) infections are usually presented as gastroenteritis in humans while in some cases it may present extra-intestinal infections and bacteremia [2–4]. Since the major reservoirs of *Salmonella enterica* for human populations are found in poultry flocks, it is expected that a decrease in *Salmonella* prevalence in poultry will also result in a decrease in the incidence of human salmonellosis [5–7]. Widespread usage of antibiotics aimed to control *Salmonella* in poultry, has led to the emergence of multiple antibiotic-resistant bacteria with many of the factors responsible for *Salmonella*s multi-drug resistance as well as multiply virulence factors are encoded by mobile genetic elements [8]. *Salmonella enterica* serovar Infantis (*S*. Infantis) is one of the most frequently isolated NTS serovar from poultry and other veterinary sources [3, 4]. Recently identified *S*. Infantis strains are characterized by the presence of mosaic megaplasmid (285 kb), termed plasmid of emerging *S*. Infantis (pESI). This conjugative plasmid contributes to virulence-associated phenotypes and pathogenicity, confers resistance to multiple antibiotics such as tetracycline and trimethoprim, heavy metals, and disinfectants [9]. This megaplasmid was also shown to be horizontally transferred to additional *salmonella* serovars as well as to other bacteria strains in the gut microbiota of infected warm-blooded hosts [10–12]. Investigation of microbial hosts and their plasmids often requires direct comparison between plasmid-containing and plasmid-free derivatives, achieved by plasmid curing. Some bacterial plasmids are extremely stable, thus plasmid elimination requires usage of curing agents or the presence of specific conditions such as thymine starvation or elevated growth temperature to increase the frequency of spontaneous segregation [13]. The usefulness of curing agents is unpredictable in many bacterial strains, as there are no standard protocols applicable to all plasmids. Another problem associated with curing procedure is the presence of plasmid addiction systems that prevent plasmid-free segregants from surviving. These systems are composed of at least two plasmid genes: one coding for a stable toxin and another for an unstable antidote (antitoxin) that can form a complex, preventing the toxin from exerting its toxicity in plasmid –containing bacteria. Plasmid loss causes release of the toxin from the existing complex, which results in growth inhibition and eventual cell death [14– 16]. The stability of pESI is achieved by the presence of at least three toxin-antitoxin addiction systems CcdAB, VapBC/VagCD, and MazEF/PemKI [17]. To prevent post-segregation killing due to the loss of pESI, we designed plasmid curing method based on specific deletion of the plasmid partition region accompanied by overproduction of specific antitoxins from a temperature sensitive plasmid in trans.

## 2. Methods

### 2.1 Bacterial strains, plasmids, and growth conditions

A field strain derived from *Salmonella enterica* serovar Infantis 119944 and containing a megaplasmid, similar to pESI # NZ_CP047882.1 [9] was isolated from poultry in Israel and used in this study (Israeli ministry of health isolate #12278). pKD46 antitoxin helper plasmid was designed in this study by cloning the sequences of *pemI, ccdA* and *vapB* antitoxins at *Nco*I restriction site of pKD46 plasmid [18]. Bacteria were grown in Luria-Bertani (LB) medium (Difco). In some cases, the medium was supplemented with ampicillin (100 or 400 μg/ml), tetracyclin (20 μg/ml) or trimethoprim (50 μg/ml).

### 2.2. pESI curing strategy

A DNA construct containing the sequences of three antitoxin genes *ccdA* (refseq NP_052634.1), *vapB* (refseq WP_001261287.1) and *pemK* (refseq WP_000288814.1), each preceded by a synthetic promoter and ending with a single stop codon, a single terminator sequence was placed after the stop codon of *pemK* (Supplementary text 1) was synthesized (GensScript; Piscatway, NJ, US). The fragment was amplified by PCR using the primers -Anti_F and Anti_R primers (Table S1) that insert a *Nco*I restriction site at both ends of the PCR product. The PCR product and a pKD46 plasmid were digested with *Nco*I and ligated using T4 DNA ligase. *S*. Infantis was transformed with the resulting plasmid (pKD46-Antitoxin), plasmid containing colonies were isolated from LB agar plates containing 400 μg/ml ampicillin. Targeted deletion of a 3 kilo base pairs (kbp) region in the pESI segregation operon was conducted in pESI and a pKD46-Antitoxin containing strain. The deletion was conducted using the lambda red recombinase procedure [18], using Inf_Seg_Del_F and Inf_Seg_Del_R primers and the pKD4 plasmid that carries a Kan^R^ cassette as the template for PCR reactions. PCR with the specified primers inserted a 50 bp stretch, homologous to the target region at each side of the Kan^R^ cassette. Cells were then transformed with the purified PCR product and grown on LB plates containing ampicillin and kanamycin. Successful deletion was verified using locus-specific primers Seg_Del_Screen_F and Seg_Del_Screen_R with the respective k2 or kt primers, and by sequencing. Kanamycin resistance transformants were grown at 25°C to eliminate pESI. This temperature was chosen to keep the temperature sensitive helper pKD46-Antitoxin plasmid. pESI curing was confirmed by loss of trimethoprim and tetracycline resistance followed by PCR using the primers, FaeH_F and FaeA_Rev for the plasmid specific type 1 fimbrial virulence gene (product size 2064bp). Positive clones were grown at 42°C to eliminate the helper pKD46-Antitoxin plasmid. At each stage, *S*. Infantis plasmid status was monitored by DNA extraction using ZR BAC DNA Miniprep Kit (Zymo Research; Irvine, CA, US) followed by agarose gel electrophoresis of plasmid linearizd with *Not*I, *Mss*I and *Xho*I restriction enzymes and a Quick-Load^®^ 1 kb Extend DNA Ladder (NEB). Plasmid curing was confirmed by loss of pESI related antibiotic resistance. To test the effect of growing temperature on pESI curing efficiency, partition region deleted bacteria were grown in LB broth for five days at 25°C and 37°C. Each day bacteria were diluted 1:1000 with fresh LB medium. At each time point the diluted culture was plated on LB agar plates, 100 colonies were transferred to fresh LB plates containing tetracycline (20 μg/ml) and the numbers of tetracycline resistant colonies were calculated.

## 3. Results

Since its discovery [3, 17], pESI and similar mega-plasmids have been extensively studied and characterized worldwide [4, 10, 12, 19, 20]. These plasmids harbor a wide array of virulence factors, multiple drug resistance genes, various mobile elements, and plasmid maintenance components. Furthermore, they possess at least three distinct toxin-antitoxin systems (MazEF/PemKI, CcdAB, and VagCD) [17], which contributes to the remarkable stability of the pESI plasmid.

To create a pESI free strain from a field strain of *Salmonella enterica* serovar Infantis isolated from chickens, our approach targeted the toxin-antitoxin systems and the partition region of this plasmid. We aimed to achieve a complete eradication of the three toxin components using one DNA construct by simultaneously overexpressing three antitoxin genes in trans, utilizing a temperature-sensitive plasmid. Through sequence analysis of the pESI plasmid (GenBank: NZ_CP047882.1), we identified the three toxin-antitoxin regions. Subsequently, the antitoxin genes *ccdA, vapB* and *pemK*, were synthesized and assembled into a single DNA sequence that incorporated the three antitoxin sequences separated by independent promoters and ending with a stop codon and a terminator sequence (Fig. 1 and supplementary text 1). This DNA construct was incorporated into a pKD46 plasmid. The pKD46 plasmid carries a heat sensitive origin of replication (repA101) and is part of the recombineering system [18] that will be used in the next step. *S*. Infantis cells were transformed with the plasmid carrying the antitoxin genes and transformants were selected and maintained using 400 μg/ml ampicillin to overcome the basal ampicillin resistance of the strain. Next, we have targeted a 3 kbp region in the mega plasmid containing multiple plasmid maintenance and plasmid partition genes including *parA* and *parM*. This region was replaced with a kanamycin resistance cassette using homologous recombination [18]. A colony with the antitoxin expressing plasmid and the partition region deletion was grown at a permissive temperature of 25°C and with 400μg/ml ampicillin to maintain the antitoxin expression plasmid for multiple generations. Resulting colonies were selected for loss of kanamycin, trimethoprim, and tetracycline resistance. In contrast to the wild type *S*. Infantis strain, pESI cured bacteria exhibited high sensitivity to tetracycline, trimethoprim, and ampicillin (Table 1). Interestingly, in contrast to the original *S*. infantis 119944 strains, our field isolate showed a high MIC of 400 μg/ml for ampicillin that was reduced to 25 μg/ml upon plasmid lost. PCR-probing with plasmid specific primers generated a single ∼2 kbp fragment in both, WT and in WT transformed with pKD46-Antitoxin plasmid strains but not in pESI cured bacteria (Fig. 2A). Plasmid extraction followed by restriction enzymes digestion resulted in formation of large fragments (>25 kbp, a collection of 7 fragments ranging from 35 to 24.9 kbp) in the WT strain, two fragments (>25 kbp and 7 kbp) in WT transformed with pKD46-Antitoxin, one fragment (7 kbp) in pESI cured strain before elimination of the helper plasmid and no fragments in the pESI and pKD46-antitoxin cured strain (Fig. 2B). To evaluate the rate of pESI lose following the deletion and overexpression of the antitoxin genes, we grew *S*. infantis containing pKD46-antitoxin and a deletion in the partition region at a permissive temperature of 25°C for four days. At time zero and every 24 h, a sample was withdrawn and plated on agar plates following with transfer of individual colonies to agar plates containing kanamycin. A fast decline in antibiotic resistance was observed with complete loss of resistance at day 4, indicating the loss of the pESI plasmid following the deletion of the partition region and overexpression of the anti-toxin genes (Fig. 3A). When the expression of the anti-toxin gene was eliminated by growing the cells at 37°C no plasmid loss was observed despite the deletion of the partition region (Fig. 3A). To eliminate the megaplasmid, it is essential to simultaneously delete the partition region and overexpress the antitoxin genes, as no tetracycline-sensitive colonies were identified following the growth of *S*. Infantise transformed with pKD46-antitoxin but without deletion of the partition region (Fig. 3B).

**Table 1.**
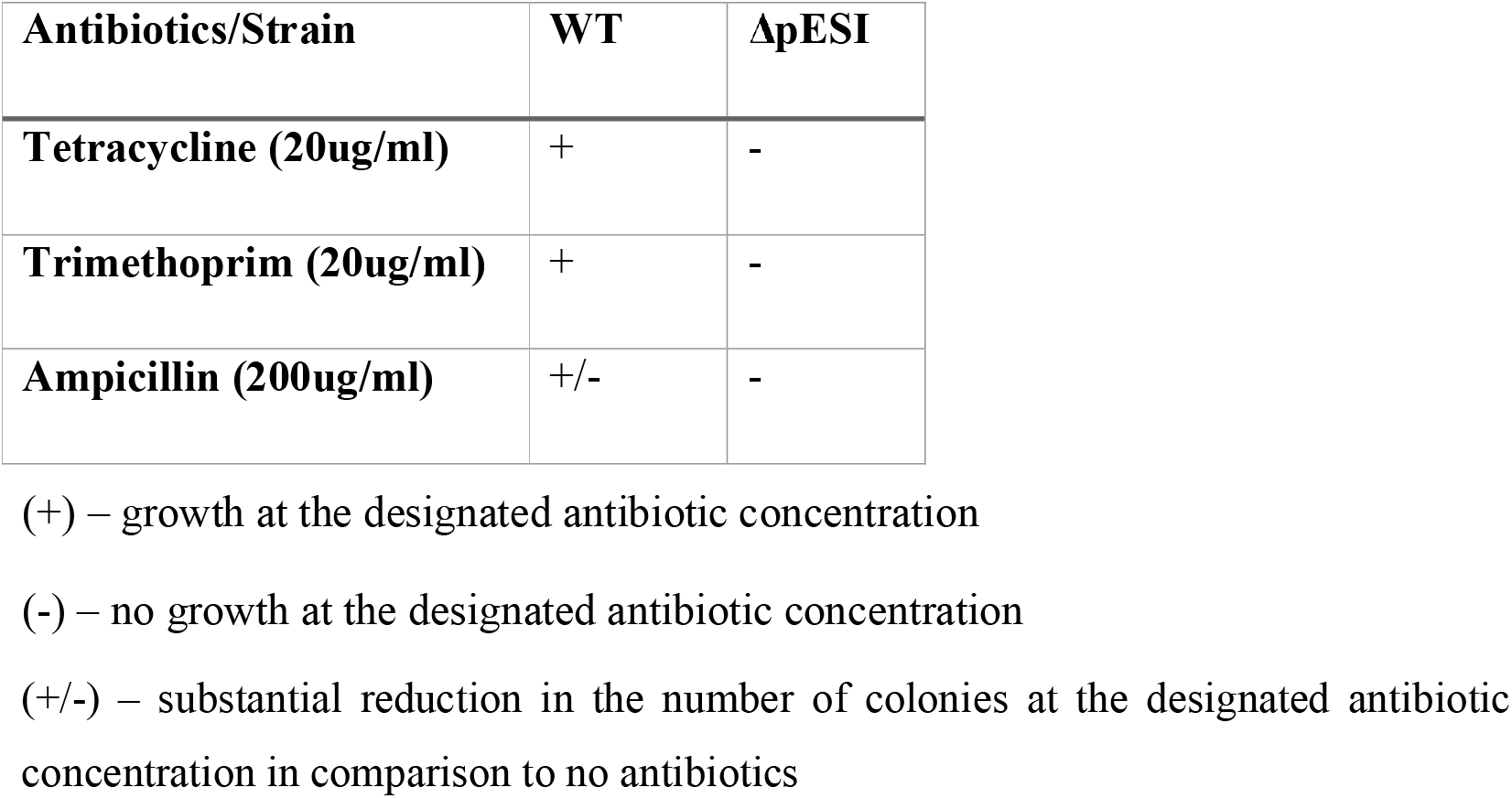
Antibiotic resistance panel of wild type and pESI cured *S*. Infantis strains.

**Fig. 1.**
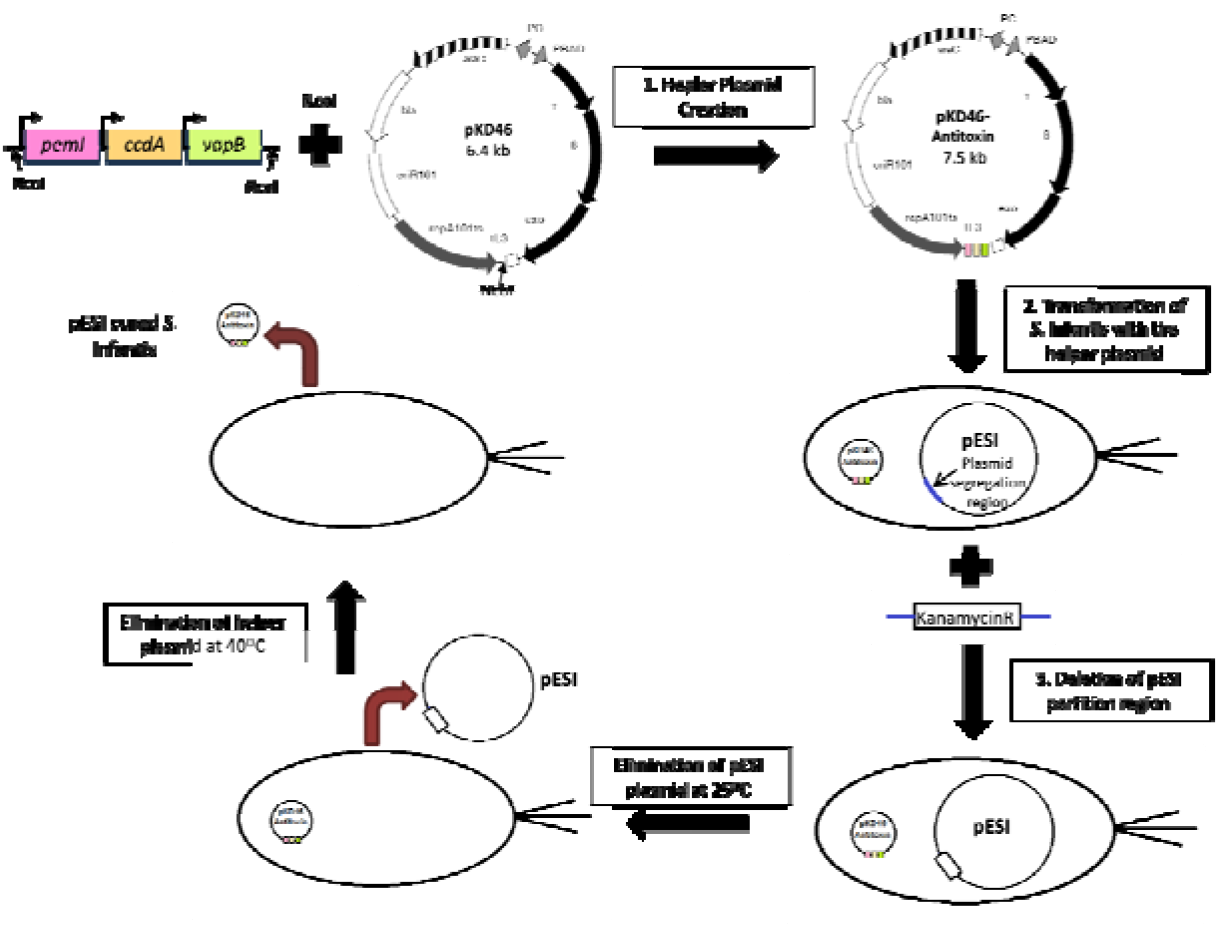
pESI curing steps. The following steps were conducted to cure *S*. Infantis strain form its pESI plasmid **1**. Construction of the helper plasmid by cloning the antitoxins genes, *pem*I, *ccd*A and *vap*B into pKD46 plasmid at *Nco*I restriction site (pKD46-antitoxin). **2**. Transformation of *S*. Infantis carrying the pESI plasmid with the pKD46–antitoxin plasmid. **3**. Deletion of pESI segregation region by □ red recombination method using KanR cassette with 50 base pairs homology to the target region. **4**. Growing of kanamycin resistant transformants at 25°C to enable curing of the pESI plasmid. **5**. Growing bacteria at 42°C to eliminate the temperature-sensitive pKD46-antitoxin plasmid.

**Fig. 2.**
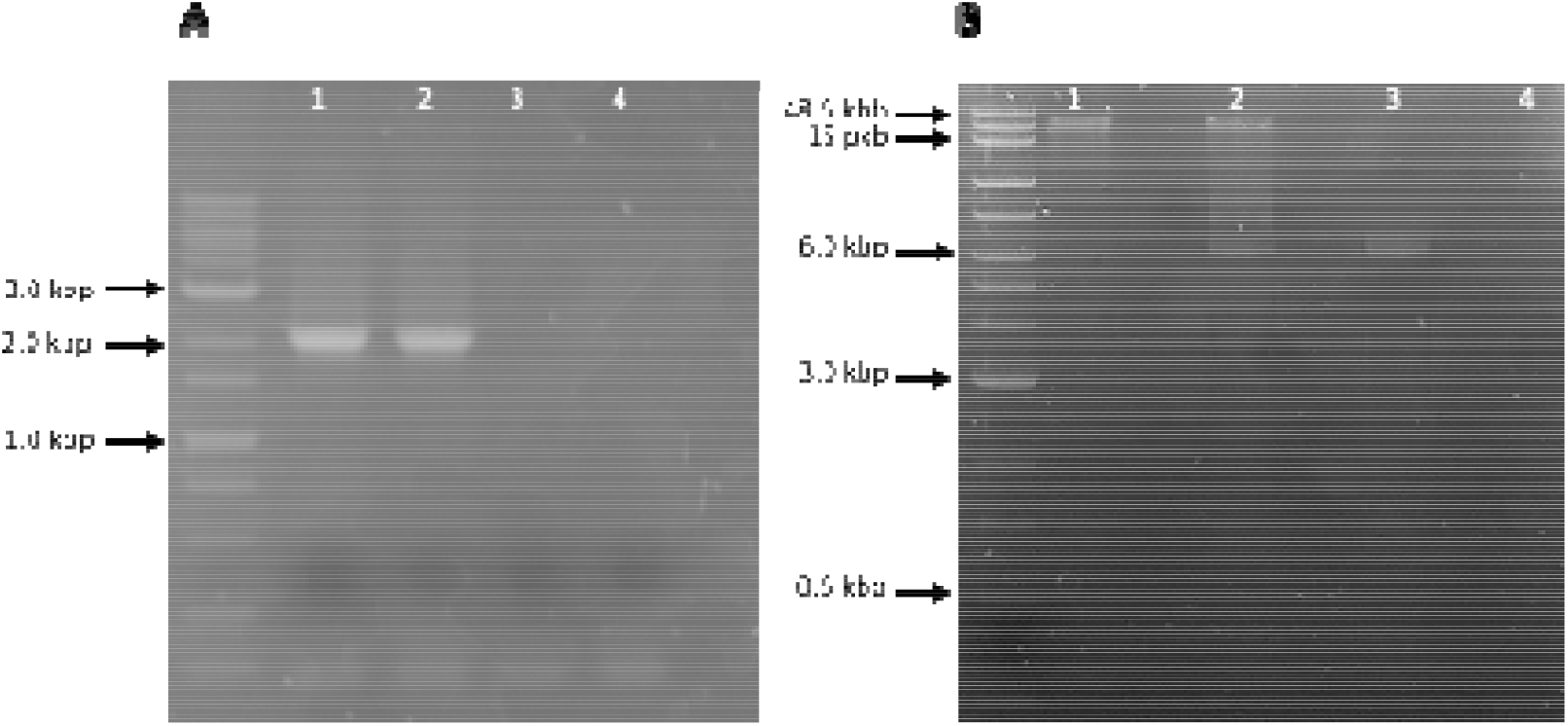
Confirmation of plasmid curing using PCR and plasmid extraction. **A**. Comparison between WT and a pESI cured strain using PCR with plasmid specific primers. Lane 1. WT strain, lane 2. WT strain transformed with pKD46-Antitoxin plasmid, lane 3. a strain following the deletion of the partition region, and Lane 4. A strain following pESI and the helper plasmid curing. **B**. Plasmids were extracted from the designated strains, and the DNA was digested with *Not*I, *Mss*I and *Xho*I restriction enzymes and analyzed using agarose gel. Lane 1. plasmids extracted from the original WT strain, lane 2. WT transformed with pKD46-Antitoxin plasmid, lane 3. WT strain transformed with pKD46-Antitoxin plasmid following growth at 25°C and lane 4. pESI cured strain following growth at 42°C.

**Fig. 3.**
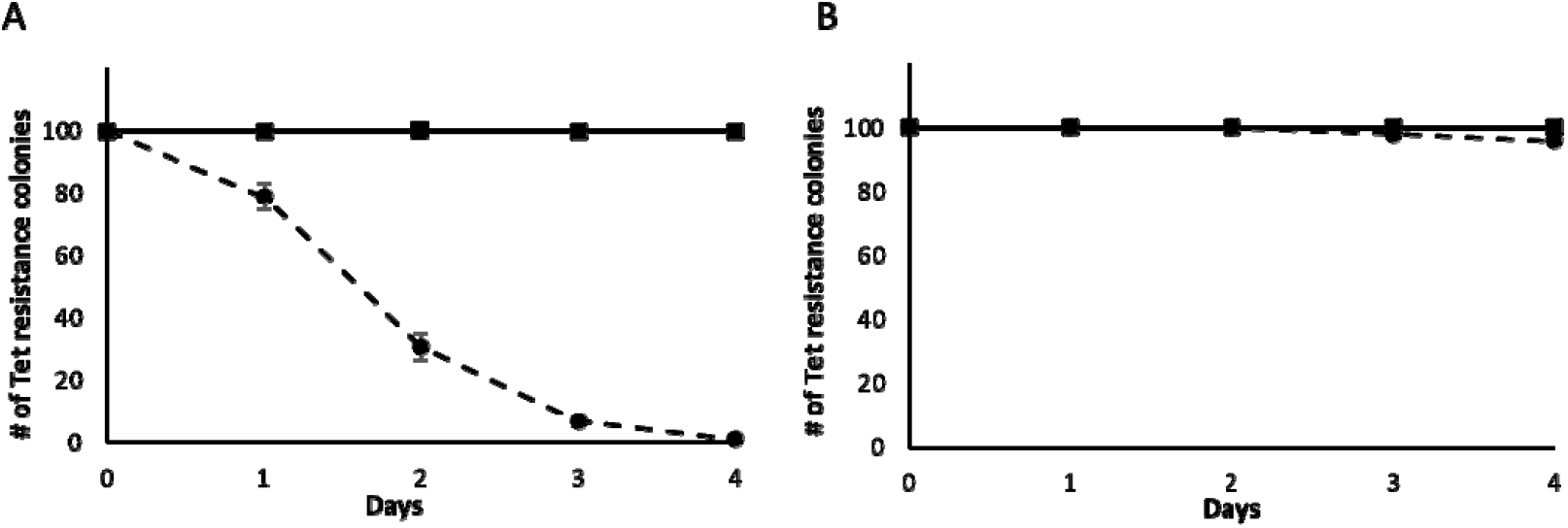
Kinetics of pESI loss following growth at 25°C and 37°C on tetracyclin containing plates. The effect of growing temperature on tetracycline resistance loss following partition region deletion and overexpression of antitoxins. At each time point a sample of the culture was withdrawn and individual colonies were grown on LB agar plates. 100 individual colonies were transferred to LB agar plates containing tetracycline and the number of resistant colonies was calculated. The data represent mean numbers from three independent experiments, each performed in duplicates. **A**. *S*. Infantis following plasmid curing steps 1 to 3 (Fig. 1), step 4 was carried in two different temperatures: 25 °C black circles and 37°C black squares. **B**. *S*. Infantis following plasmid curing steps 1 to 2 (Fig. 1), step 4 was carried in two different temperatures: 25 °C black circles and 37°C black squares.

## 4. Discussion

Natural plasmids in *Salmonella enterica* encode various functions involved in virulence and persistence of the pathogen [21, 22]. Elucidation of bacterial phenotypes associated with these traits, requires efficient plasmid curing. Various methods have been used to generate plasmid-less strains of *Salmonella enterica* [23]. However, most of them either have low efficiency or require cultivation under stressful conditions promoting the accumulation of second-site mutations [24]. Major improvements to those methods were introduced following the availability of plasmid sequences, recognition of PSK mechanisms, their integration into curing systems, and the development of novel DNA editing tools [25–27]. For instance, Hale et al. developed a highly efficient curing system that targets both the PSK systems and the replication mechanism, incorporating them into a single curing plasmid. These inhibition of the replication mechanism and over-expression of the anti-toxin genes resulted in 99% loss of plasmid within one day as demonstrated for curing of *E. coli* O157:H7 from the pO157 plasmid [26]. Combining this system with a self-transmissible plasmid demonstrated the ability of this system to eliminate plasmids in-vivo, highlighting its potential in combating multiple drug resistance [26, 28, 29]. We developed a parallel, simple, and rapid method for *in vitro* curing of *S*. Infantis from its pESI megaplasmid based on widely available and validated molecular tools to target the PSK systems by over expression of the anti-toxin genes and by physical deletion of the partition mechanism. Targeting the partitioning genes in *S*. infantis, resulted in curing of plasmids in over 99% of the population in 4 days (Fig. 3A). Direct deletion of large regions within the plasmid can prove useful in cases where the partition, replication, or incompatibility mechanisms are not fully characterized, or when the construction and transformation of large DNA fragments are limiting factors. This method could also be useful for curing a particular plasmid of known sequence from other bacteria. Plasmid-free bacteria generated by this method can be utilized both in applied biotechnology such as generation of veterinary vaccine strains with minimized risks of inducing mutations [30] as well as in basic research, providing theoretical data regarding bacterial virulence in the case of plasmid mediated pathogenicity and antibiotic resistance.

## Abbreviations

pESI: plasmid of emerging *Salmonella* Infantis;
PSKs: post-segregational killing systems;
NTS: non-typhoid *Salmonella*;
MIC: minimal inhibitory concentration.

## Author contributions

CK and NG, Conceptualization; NG, CK and IY, Data curation; CK and IY, Funding acquisition; NG, Investigation; NG and IY, Writing, review & editing.

## Supplementary materials

**Table S1.**
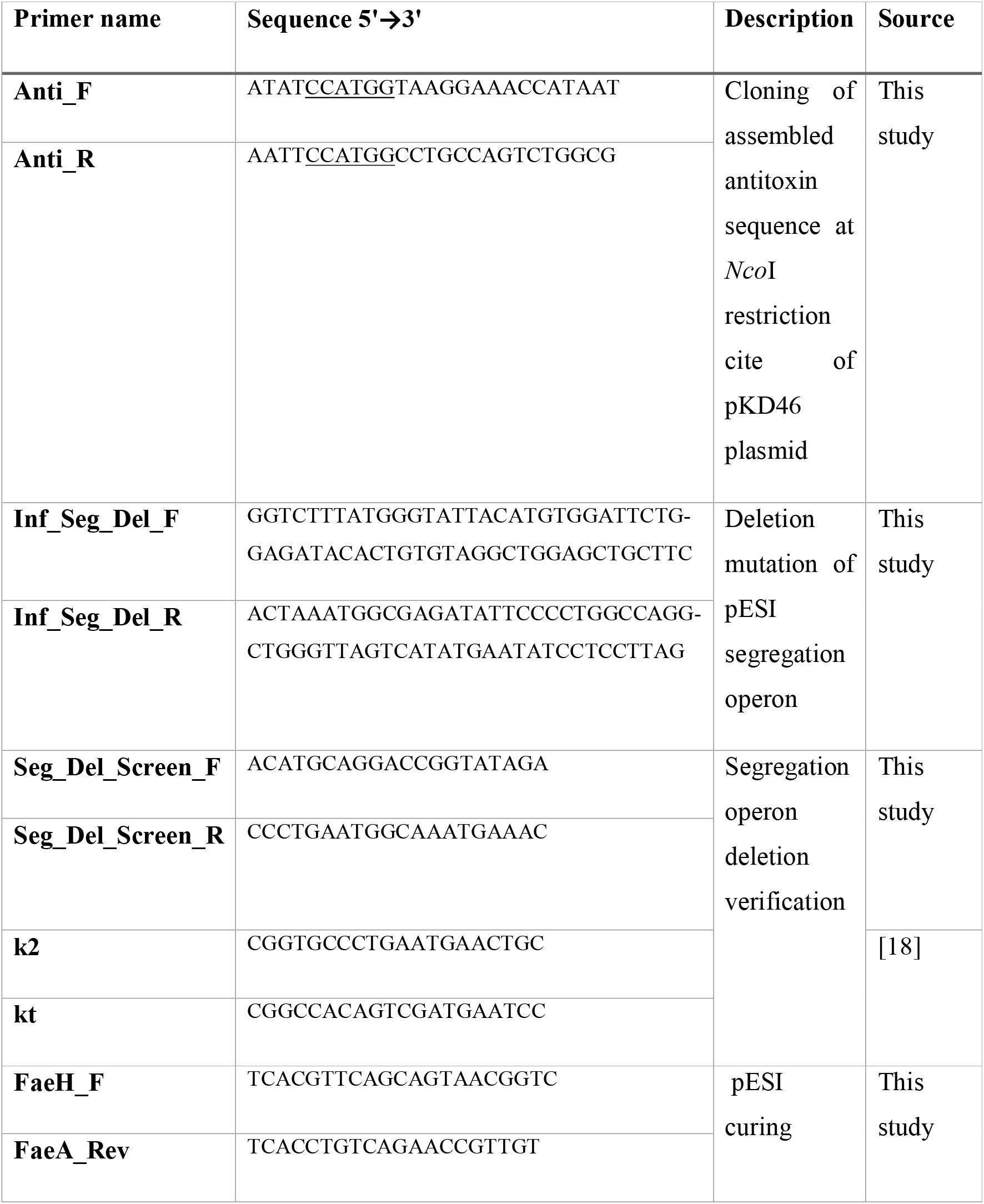
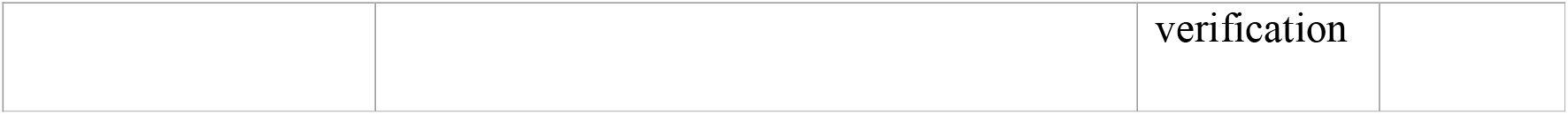
Primers used in this study.

## Supplementary text 1

Sequence of the synthetic antitoxin expression cassette

`>pemK_ccdA_vapB`

~~~
ccatggtaaggaaaccataatgaatctgactaaaaattgcgctccgccagccattga
tgttatattaaaatataatatccggaggtgctct**ATG**TATACCACTCGTCTGAAAAA
GGTTGGCGGATCCGTCATGCTGGCGGTTCCTCCCGCCGTGTTGAAAACGCTGGCTCT
GTCGACAGACAGCGAAGTGGGAATGACCATTGATAATGGCTGCCTGATTATTGAACC
CCAGAAACGGCCCCGTTATTCGCTTGAGGAACTGCTGGCACAGTGCGATCCGCACGC
CGAAATGAGCGATGAGGATAGGGAATGGATGGATGCGCCTGCAGTGGGTAAGGAGAT
CCTG**TAA**taagaaaacatcacggcagggaaaatcgtattaaaaaacttatggcgaat
taatggtatgctttttctggctatcaccaaggtgtgtctttcatctatactgatagg
catattataggtatgttaatgagggcttatc**ATG**AAGCAGCGTATTACTGTCACCAT
AGACAGTGACAGCTATCAGTTACTCAAGTCCGCAGACGTGAATATCTCCGGCCTGGT
AAATACTGCCATGCAGAAGGAGGCCCGCCGTCTGCGTGCGGAACGCTGGCAGGCGGA
AAATCAGGAAGGGATGGCTGAGGTCGCCCGGTTTATTGAAGCGAACGGCTCCTTTGC
AGACGAAAACAGGGACTGG**TGA**catatcctcaccatctgcatatgctgcatatatac
atatgctgaacatctgcctatactgaacagatgcacacacccggtgcaatgaacatc
attccggatatgcatcacatatccacacttacccgggagagaatc**ATG**AGAACCGTA
TCTATTTTTAAAAACGGCAACAACCGCGCCATCCGCCTTCCCCGCGATCTGGATTTT
GATGGAGTGAGCGAGCTGGAAATCGTCCGGGAAGGGGACAGCATCATCCTGCGCCCT
GTCCGGCCGACCTGGGGCTCGTTCGCGCAGCTCGAAAAAGCTGACCCGGGCTTTATG
GCGGAGCGTGAGGACGTTGTCAGCGACGAAGGACGATTTAACCTG**TGA**gtatccgga
gtttctggcgccgctccgccagactggcaggccatgg
~~~

Uppercase letters – antitoxins open reading frame.

Lowercase letters – promoter region.

Lowercase underlined letters – terminator region.

ORF1 - PemK, ORF2 – CcdA, ORF3 – VapB

